# Adipose-tissue derived signals control bone remodelling

**DOI:** 10.1101/2020.03.02.972711

**Authors:** Maria-Bernadette Madel, He Fu, Dominique D. Pierroz, Mariano Schiffrin, Carine Winkler, Anne Wilson, Cécile Pochon, Barbara Toffoli, Jean-Yves Jouzeau, Federica Gilardi, Serge Ferrari, Nicolas Bonnet, Claudine Blin-Wakkach, Béatrice Desvergne, David Moulin

## Abstract

Long bones from mammals host blood cell formation and contain multiple cell types, including adipocytes. Physiological functions of bone marrow adipocytes are poorly documented. Herein, we used adipocyte-deficient PPARγ-whole body null mice to investigate the consequence of total adipocyte deficiency on bone homeostasis in mice. We first highlight the dual bone phenotype of PPARγ null mice: on the one hand the increase bone formation and subsequent trabecularization extending in the long bone diaphysis, due to the well-known impact of PPARγ deficiency on osteoblasts formation and activity; on the other hand, an increased osteoclastogenesis in the cortical bone. We then further explore the cause of this unexpected increased osteoclastogenesis using two independent models of lipoatrophy, which recapitulated this phenotype. This demonstrates that hyperosteoclastogenesis is not intrinsically linked to PPARγ deficiency, but is a consequence of the total lipodystrophy. We further showed that adiponectin, a cytokine produced by adipocytes and mesenchymal stromal cells is a potent inhibitor of osteoclastogenesis *in vitro* and *in vivo*. Moreover, pharmacological activation of adiponectin receptors by the synthetic agonist AdipoRon inhibits mature osteoclast activity both in mouse and human cells by blocking podosome formation through AMPK activation. Finally, we demonstrated that AdipoRon treatment blocks bone erosion in vivo in a murine model of inflammatory bone loss, providing potential new approaches to treat osteoporosis.

## Introduction

Bone homeostasis is a result of constant remodelling activities, with a balance of bone resorption and bone formation. The most prevalent disorder of bone homeostasis is osteoporosis, affecting men and women, with a particular high prevalence in post-menopausal women. The low efficacy of actual anti-osteoporotic therapeutic options highlights the need of better understanding the pathophysiological processes at work.

The role of adipocyte in bone homeostasis is one of the factors raising great interest, but it faces quite some complexity. An illustration of this complexity, is the fact that obesity is traditionally considered to be protective against osteoporosis, consequently to an increased proliferation and differentiation of osteoblasts and osteocytes stimulated by mechanical load (Felson et al., 1993). However, obesity in postmenopausal women and in men correlates with increased marrow adiposity, alteration of the bone microarchitecture, and increased risk of fracture (Sheu and Cauley, 2011).Thus, the respective roles of ageing, increased BMI, and visceral *vs* bone marrow fat on bone homeostasis remain difficult to disentangle (Cornish et al., 2018; Horowitz et al., 2017).

Peroxisome proliferator-activated receptors (PPARs) are ligand-activated transcription factors that belong to the nuclear hormone receptor superfamily. PPARγ is a key factor and master regulator in adipocytes development and functions (Tontonoz and Spiegelman, 2008). Indeed, loss-of-function studies both *in vitro* and *in vivo* have clearly demonstrated that PPARγ is essential for the formation of adipose tissue (Barak et al., 1999; Rosen et al., 1999; Wang et al., 2013) and mature adipocyte homeostasis (Imai et al., 2004). Recently, we demonstrated that mice carrying a full body deletion of PPARγ (*Pparg*^*Δ/Δ*^) are totally devoid of adipocytes (Sardella et al., 2018). The activity of PPARγ is also central in bone homeostasis by modulating both bone formation by osteoblasts and bone resorption due to osteoclast activity. First, adipocytes and osteoblasts share same mesenchymal progenitors, and the cell fate decision between adipocyte and osteoblast lineages depends on various signals (Calo et al., 2010; Idris et al., 2009; Kawai and Rosen, 2010; Wan, 2010). Activation of PPARγ inhibits osteoblast formation, and thus bone formation, by favouring adipocyte differentiation (Ali et al., 2005; Rzonca et al., 2004). Along this line, *in vivo* evidence indicates that mice heterozygous for a null allele of the gene encoding PPARγ (*Pparg*) exhibit increased osteoblastogenesis, which results in increased bone mass with a doubling of the rate of new bone formation, when compared to control mice. Moreover, PPARγ also directly inhibits osteoblast activity, independently of adipogenesis, as demonstrated by the consequence of osteoblast-specific *Pparg* deletion in osteoblast differentiation and trabecular bone formation (Sun et al., 2013). Second, in bone resorption, PPARγ promotes osteoclastogenesis, as illustrated by increased osteoclast numbers in aged mice treated with thiazolidinediones (Lazarenko et al., 2007). Conversely, mice in which *Pparg* is specifically deleted in endothelial and hematopoietic cells, (i.e., affecting osteoclast progenitors but not osteoblasts), exhibit decreased osteoclastogenesis compared to wild-type mice, provoking osteopetrosis with impaired bone resorption and reduced medullary cavities (Wan et al., 2007). However, a recent controversy on the role of PPARγ in osteoclast was raised by Zou et al. who found that only the pharmacological, and not the physiological activation of PPARγ affects osteoclastogenesis (Zou et al., 2016). Finally, hyperactivation of PPARγ via agonist treatment causes accumulation of adipocytes in the bone marrow in thiazolidinedione-treated rodents, whereas increased bone loss and bone fractures have been correlated with thiazolidinedione therapy in humans (Grey, 2008; Kahn et al., 2008; Lazarenko et al., 2007).

Herein, we used mice carrying a PPARγ full body deletion to explore the consequences of the lack of adipocyte on bone homeostasis, and demonstrate that as expected, there is a strong increased bone density of the trabecular bone. However, this phenotype is contrasted by an important osteoclastogenesis. Using two other complementary models of genetically-induced lipoatrophic mice, i.e. mice carrying an adipose-tissue specific deletion of PPARγ and AZIP mice, we then further explore the cross-talk between adipocytes and osteoclasts, whose alteration leads to hyperosteoclastogenesis and osteoporosis.

## Results

In order to evaluate the impact of adipose tissue on bone homeostasis, we studied lipoatrophic mice that we recently generated. These mice, hereby called *Pparg*^*Δ/Δ*^ carry a whole body deletion of PPARγ and were obtained through epiblastic (Sox2 promoter) *cre* deletion of the Pparγ allele (Sardella et al., 2018). Bone analyses of males and females one-year-old mice first show that these mice displayed dramatic alterations of both trabecular and cortical bones (Fig 1, supplementary Fig S1). The three-dimensional images generated by trabecular bone microcomputed tomography (3D micro-CT) reconstruction of the femurs of one-year old male and female mice highlighted a remarkably high trabecularization of the bone architecture in *Pparg*^*Δ/Δ*^ mice, which extended along the shaft and almost reached the mid-diaphysis (VIDEO 1, Fig. 1a, Table 1a, supplementary Fig. S1). In contrast, the cortical bone was found to be extremely porous at the epiphysis and the diaphysis. Owing to hypertrabecularization and high cortical porosity, the frontier between these two types of bone along the diaphysis remained difficult to establish.

**Table 1.** **A)** Micro-CT data were obtained from various bones in *Pparg*^*fl/+*^ and *Pparg*^*Δ/Δ*^ of 1 year-old female mice. **B)** Similar evaluation of bones from 3 month- and 5/6 month-old female mice. Values are given as mean ± standard deviation. TV, total volume; BV, bone volume; Tb N, trabecular number; Tb Th, trabecular thickness; Tb Sp, trabecular separation. ^*^: p <0.05 vs *Pparg*^*fl/+*^.

| **1 year** | ***Pparg*<sup>fl/+</sup>** | ***Pparg*<sup>Δ/Δ</sup>** |
| --- | --- | --- |
| *No. of animals assessed* | 15 | 14 |
| Cortical TV (mm<sup>3</sup>) | 1.498 ± 0.023 | 1.327 ± 0.031* |
| Cortical BV (mm<sup>3</sup>) | 0.723 ± 0.012 | 0.667 ± 0.028* |
| Cortical thickness (μm) | 241.8 ± 4.5 | 208.0 ± 10.4* |
| Dist Femoral BV/TV (%) | 3.41 ± 0.49 | 14.38 ± 2.26* |
| Dist Femoral conn density (mm<sup>3</sup>) | 9.18 ± 2.57 | 81.89 ± 13.62* |
| Dist Femoral Tb N (/mm) | 2.39 ± 0.09 | 3.61 ± 0.22* |
| Dist Femoral Tb Th (μm) | 54.37 ± 2.48 | 57.28 ± 3.52* |
| Dist Femoral Tb Sp (μm) | 431.34 ± 18.4 | 291.10 ± 9.70* |
| Vertebral BV/TV (%) | 17.33 ± 1.38 | 29.57 ± 3.30* |
| Vertebral conn density (mm<sup>3</sup>) | 127.02 ± 14.89 | 519.44 ± 73.33* |
| Vertebral Tb N (/mm) | 3.42 ± 0.17 | 6.29 ± 0.53* |
| Vertebral Tb Th (μm) | 51.43 ± 1.88 | 47.04 ± 2.30 |
| Vertebral Tb Sp (μm) | 311.13 ± 16.23 | 177.30 ± 15.73* |

Table 1. **A)** Micro-CT data were obtained from various bones in *Pparg*<sup>fl/+</sup> and *Pparg*<sup>Δ/Δ</sup> of 1 year-old female mice.
| **3 months** | ***Pparg*<sup>fl/+</sup>** | ***Pparg*<sup>Δ/Δ</sup>** |
| --- | --- | --- |
| *No. of animals assessed* | 3 | 6 |
| Cortical TV (mm<sup>3</sup>) | 0.585 ± 0.038 | 0.562 ± 0.019 |
| Cortical BV (mm<sup>3</sup>) | 0.261 ± 0.016 | 0.266 ± 0.011 |
| Cortical thickness (μm) | 189.7 ± 5.2 | 197.7 ± 5.6 |
| Dist Femoral BV/TV (%) | 9.82 ± 0.79 | 9.88 ± 1.50 |
| Dist Femoral conn density (mm<sup>3</sup>) | 88.63 ± 14.21 | 141.96 ± 22.10 |
| Dist Femoral Tb N (/mm) | 4.28 ± 0.09 | 4.58 ± 0.19 |
| Dist Femoral Tb Th (μm) | 48.67 ± 2.22 | 39.93 ± 2.38* |
| Dist Femoral Tb Sp (μm) | 232.70 ± 5.19 | 215.98 ± 10.35 |
| **5-6 months** |  |  |
| *No. of animals assessed* | 8 | 10 |
| Cortical TV (mm<sup>3</sup>) | 1.18 ± 0.05 | 1.18 ± 0.04 |
| Cortical BV (mm<sup>3</sup>) | 0.0541 ± 0.023 | 0.0548 ± 0.014 |
| Cortical thickness (μm) | 203.75 ± 4.80 | 202.80 ± 5.33 |
| Dist Femoral BV/TV (%) | 7.78 ± 1.17 | 8.55 ± 1.07 |
| Dist Femoral conn density (mm<sup>3</sup>) | 38.90 ± 8.70 | 84.30 ± 14.85* |
| Dist Femoral Tb N (/mm) | 3.13 ± 0.13 | 3.75 ± 0.22* |
| Dist Femoral Tb Th (μm) | 53.01 ± 1.62 | 45.93 ± 2.24* |
| Dist Femoral Tb Sp (μm) | 327.46 ± 15.03 | 283.59 ± 21.33 |
| Vertebral BV/TV (%) | 15.49 ± 1.31 | 21.44 ± 1.79* |
| Vertebral conn density (mm<sup>3</sup>) | 112.73 ± 12.56 | 275.53 ± 33.10* |
| Vertebral Tb N (/mm) | 3.71 ± 0.13 | 4.76 ± 0.17* |
| Vertebral Tb Th (μm) | 49.03 ± 1.17 | 46.39 ± 1.14 |
| Vertebral Tb Sp (μm) | 278.37 ± 10.66 | 215.1 ± 8.32* |

**Figure 1:**
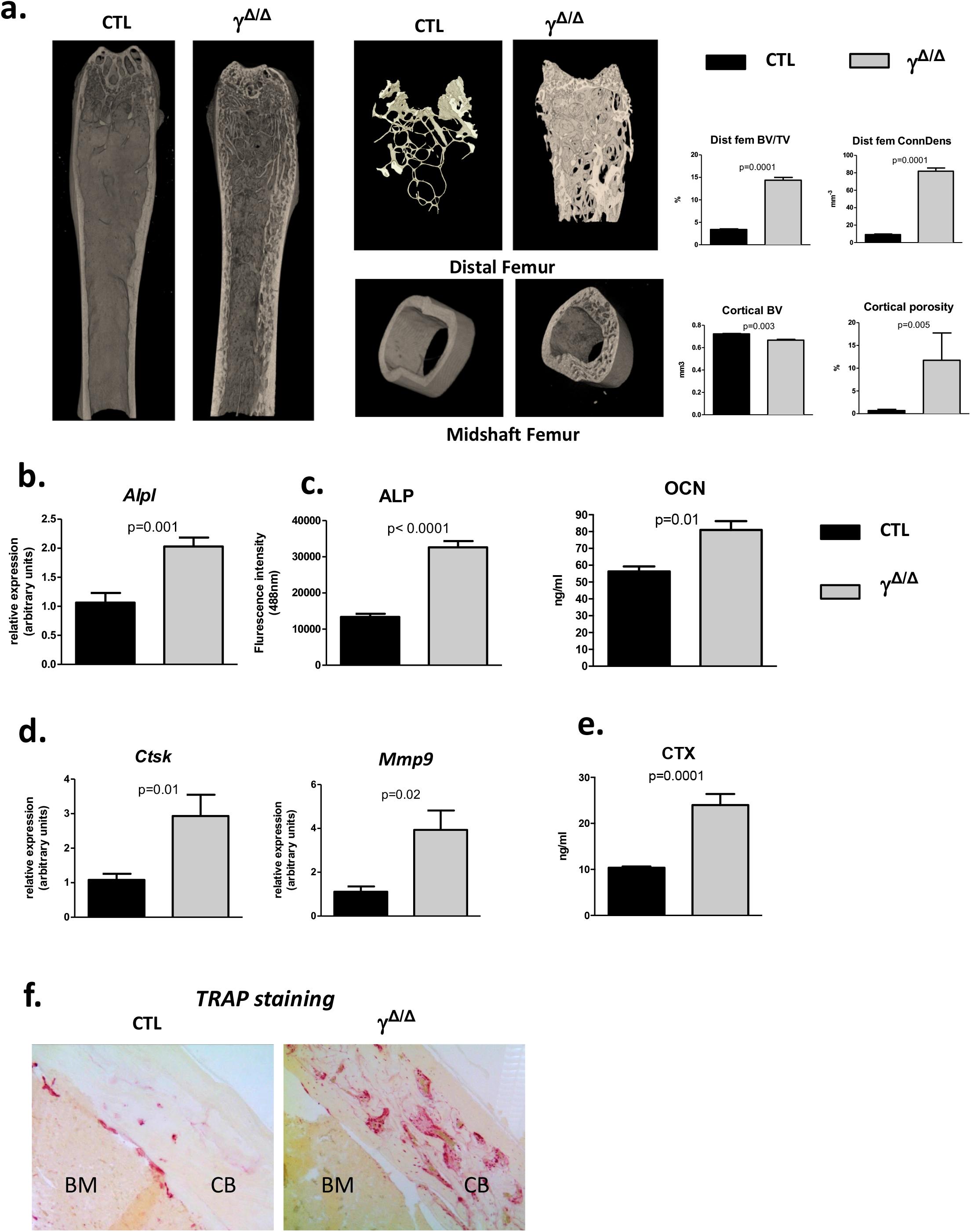
Epiblastic PPARγ-deletion induces trabecular and cortical bone alteration. **a.** Micro-CT analysis of femur from *Pparg*^*Δ/Δ*^ (γ^*Δ/Δ*^) one-year old female mice or control female littermates (CTL). Histograms on the right show for the different genotypes (black bars for CTL and grey bars for γ^*Δ/Δ*^) the percentage bone volume/trabecular volume (BV/TV) in % and connectivity density (ConnDens) of distal femoral trabecular bone (upper panels), and the cortical bone volume (BV) in mm^3^ and the percentage of cortical porosity of femur (lower panels); n = 5 WT and 5 γ^*Δ/Δ*^. All mice were 1-year-old. **b**. RT-qPCR analysis of Alkaline phosphatase (*AlpI*) gene expression in femur from CTL and γ^*Δ/Δ*^; n = 6 WT and 7 γ^*Δ/Δ*^. **c.** ELISA measurements of serum Alkaline phosphatase (ALP) and osteocalcin (OCN); n = 8 CTL and 8 γ^*Δ/Δ*^. **d.** RT-qPCR analysis of cathepsin K (*Ctsk*) and *Mmp9* expression in femur from CTL and γ^*Δ/Δ*^; n = 7 CTL and 7 γ^*Δ/Δ*^. **e.** ELISA measurements of serum Carboxy-terminal Collagen I telopeptide (CTX); n= 8 CTL and 8 γ^*Δ/Δ*^. **f.** Representative images of TRAcP staining of decalcified sections of femurs, at mid-shaft levels, from control (CTL) or *Pparg*^*Δ/Δ*^ (γ^*Δ/Δ*^) mice. CB: cortical bone; BM: bone marrow. Data are presented as mean ± S.E.M. Statistical significance was determined by two-tailed unpaired *t*-test.

The phenotype being more marked in female (supplementary Fig S1), we further evaluated in female whether the bone alterations appear during development or appear progressively with age, by performing micro CT measurements at 3, 5 and 6 months. A quantification of the diverse parameters is shown in Table 1b. More particularly, we found that the increased trabecular number and connectivity density in both vertebral bodies and distal femoral metaphysic bones appeared at 5-6 months of age (Table 1).

Molecular analyses confirmed an increased trabecular bone formation, as exemplified by increased levels of alkaline phosphatase mRNA (*Alp1*) in the bone, evaluated by RT-qPCR and increased levels of alkaline phosphatase (ALP) and osteocalcin (OCN) in the serum measured by ELISA (Fig.1b and c). To evaluate whether osteocytes, which come from osteoblasts and are mature permanent bone cells, *versus* osteoblast populations were altered in these mice, we measured the mRNA levels of sclerostin (*Sost*) and keratocan (*Kera*), which are expressed by osteocytes and osteoblasts, respectively (Jilka et al., 2014). Both markers were unchanged (supplementary Fig. S2). Three-D micro-CT of the fourth lumbar vertebrae clearly demonstrated increased trabecular bone volume fraction and trabecular connectivity density, indicating increased osteoblastogenesis in *Pparg*^*Δ/Δ*^ (supplementary Fig. S2), in agreement with previous report (Akune et al., 2004). These results are consistent with the pivotal role of PPARγ on the fate of bone marrow mesenchymal stromal cells, with previous reports showing increased osteoblastogenesis when PPARγ signalling is pharmacologically or genetically blocked (Takada et al., 2009).

Remarkably, the increased expression in the long bones of *Pparg*^*Δ/Δ*^ mice compared to control mice of bone resorbing markers such as cathepsin K (*Ctsk*) and *Mmp9*, which is a key proteinase involved in the recruitment and activity of osteoclasts and endothelial cells for the development of long bones, particularly in the diaphysis core (Engsig et al., 2000; Jemtland et al., 1998), (Fig. 1d) suggest a parallel increase in osteoclastogenesis. This is further supported by a parallel augmentation of serum levels of carboxy-terminal collagen I telopeptide (CTX) (Fig. 1e). Indeed, TRAcP staining on femoral sections (Fig. 1f) confirmed the increased presence of bone-resorbing osteoclasts in cortical bone, explaining the observed cortical porosity.

PPARγ is known to play a role in activating osteoclastogenesis. More specifically, Tie2^Cre^-directed deletion of PPARγ severely impaired osteoclastogenesis, whereas PPARγ synthetic agonists exacerbated it (Wan et al., 2007). This role of PPARγ in osteoclast has been recently questioned by the study of Zou *et al.*, showing that the *in vivo* effect of PPARγ on osteoclast formation is only seen upon pharmacological activation of PPARγ but neither on physiological nor on pathological osteoclast formation (Zou et al., 2016). This raised the question whether the elevated osteoclast activity observed in cortical bone is attributable to lipoatrophy, or to unforeseen intrinsic effects of PPARγ deficiency in the osteoclast lineage. To disentangle these two possibilities, we used another PPARγ-independent lipodystrophic mouse model. AZIP^tg/+^ mouse is a hemizygote transgenic mouse strain in which a dominant negative protein of key B-ZIP transcription factors (A-ZIP) blocks adipocyte development. AZIP^tg/+^ mice are born with no white adipose tissue and markedly decreased brown adipose tissue (Moitra et al., 1998). In support of the lipoatrophy hypothesis, micro-CT analyses of femur of AZIP^tg/+^ mice revealed an increased cortical porosity (Fig 2a). Finally, we generated lipoatrophic mice with wild-type hematopoietic lineage by invalidating PPARγ in adipose tissue (Adipoq^cre^ PPARγ fl/fl, γ^*FatKO*^ (Wang et al., 2013), called herein PPARγ^adipoKO^). TRAcP staining of cortical bone and micro-CT analysis in these mice revealed increased osteoclast numbers and activity with an enhanced cortical porosity, as observed in *Pparg*^*Δ/Δ*^ mice (Fig. 2a-c and supplementary Fig. S3). These two independent mouse models confirmed that adipocyte deficiency is pivotal to the cortical bone phenotype and the increased osteoclast number observed here.

**Figure 2:**
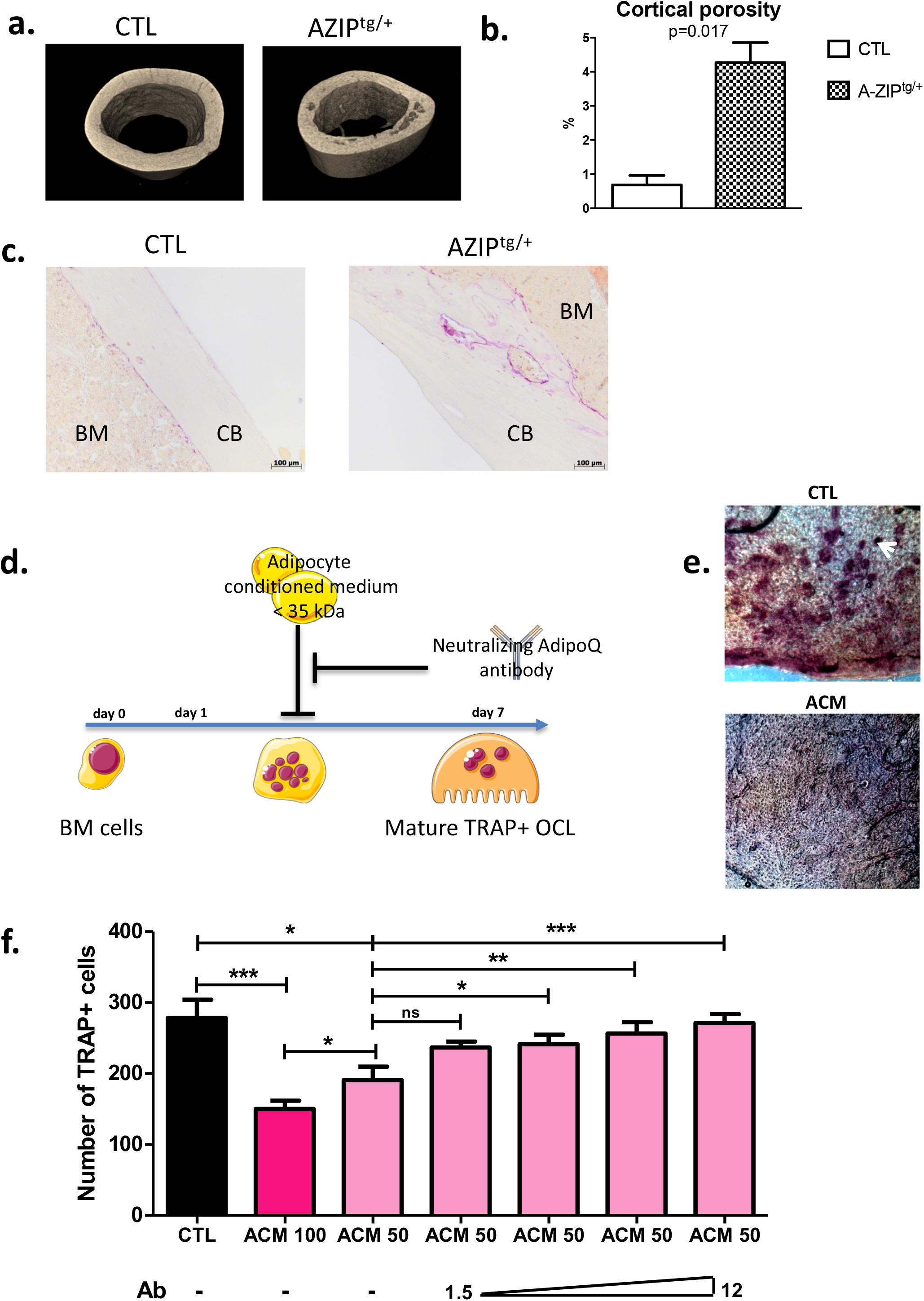
Lipoatrophy is responsible for unregulated osteoclastogenesis. **a.**. Representative 3D sections of midshaft femur from one year-old female A-ZIP^tg/+^ (n=6) and CTL (n=6) littermates. **b.** Femoral cortical porosity measured by micro-CT analysis of femur from A-ZIP^tg/+^ (n=6) and control (CTL; n=6) littermates. **c.** Representative images of TRAcP staining of longitudinal sections of decalcified femurs, at mid-shaft levels, from control (CTL) or A-ZIP^tg/+^ mice; CB: cortical bone; BM: bone marrow. **d**. Experimental design of the *in vitro* assay used to test anti-osteoclastogenic activity from adipocyte secretome. **e**. Calvaria resorption pit assay. Representative sections of skulls from 14-day-old wild-type pups cultured for 2 weeks in complete αMEM in the presence of conditioned medium from either undifferentiated 3T3-L1 preadipocytes (CTL, n=3) or mature 3T3-L1 adipocytes (ACM, n=3), after TRAP staining. Top panel: the white arrow indicates one example of resorption pit seen in many places on the skull with CTL medium. **f**. One day after plating, wild type BM cells were cultured for 6 more days in the presence of M-CSF and RANKL (osteoclast differentiation media) in addition to conditioned medium prepared from differentiated (ACM: adipocyte-conditioned medium) or undifferentiated (CTL: control-conditioned medium) 3T3-L1 preadipocytes. Different doses of a specific blocking antibody against adiponectin (1.5–12.0 μg/ml) were added to the conditioned medium (ACM50) during osteoclast differentiation (ACM 100 means pure ACM; ACM50 means 1:1 diluted ACM with osteoclast differentiation medium).

One consequence of lipodystrophy observed in all three models is a severe type 2 diabetes with signs of diabetic nephropathy (Moitra et al., 1998; Toffoli et al., 2017; Wang et al., 2013). Given that chronic kidney diseases are often accompanied by hyperparathyroidism and result in complex, combined catabolic and anabolic activities in the bones (Levin et al., 2007), we evaluated parathyroid hormone (PTH) levels. As shown in supplementary Figure S4, no statistically significant differences were observed between *Pparg*^*Δ/Δ*^ mice and their control littermates in either the circulating levels of PTH or the mRNA levels of the PTH receptor Pth1r in the bones or kidneys. Notably, serum PTH levels were also unchanged in AZIP^tg/+^ mice (supplementary Fig S4), suggesting that the PTH pathway is unlikely to support the increased bone remodelling observed in lipodystrophic mice.

To investigate how adipocytes regulate osteoclastogenesis, and to evaluate whether factors produced by adipocytes within the bone marrow can influence osteoclastogenesis, we performed an osteoclastogenesis assay using bone marrow cells from wild-type mice cultured in the presence of either control medium or adipocyte-conditioned medium (ACM in figure 2). Adipocyte-conditioned medium was isolated from culture of adipocyte-differentiated 3T3-L1 cells whereas control medium (CTL in figure 2) was obtained from cultures of undifferentiated pre-adipocytes 3T3-L1 cells. As seen in Figure 2, the presence in bone marrow cells culture of adipocyte-conditioned medium, compared to pre-adipocyte-conditioned medium, provoked a substantial decrease in the number of mature osteoclasts. Consistently, adipocyte-conditioned medium significantly inhibited bone resorption as assessed by the decrease number of resorption pits in the calvaria resorption assay (Fig. 2d-f). These data demonstrate that adipocyte-secreted mediators are responsible for the inhibition of RANKL-induced osteoclast differentiation *in vitro*.

Next, we searched for component(s) of ACM responsible for this anti-osteoclastogenic activity. Interestingly, the ACM inhibitory activity was maintained after filtering out all components larger than 35 kDa from the conditioned medium. The molecular weights of the major adipokines, adiponectin and leptin, are below 35 kDa and their circulating levels are not or barely detectable in *Pparg*^*Δ/Δ*^ mice, as we previously demonstrated (Gilardi et al., 2019). Unlike *Leptin*, *Adiponectin* is preferentially expressed by adipocyte and by bone-marrow MSC, and is a well-characterized target gene of PPARγ (Iwaki et al., 2003). Western-blot analyses confirmed that adiponectin was present in large amounts in ACM but not in CTL medium (supplementary Fig. S5). We therefore tested whether blocking adiponectin would affect ACM-mediated inhibition of osteoclastogenesis *in vitro*. As shown in Figure 2f, the addition of a blocking adiponectin antibody reversed the reduction in osteoclast number in a dose-dependent manner. These results support that the soluble factors secreted by adipocytes, in particular adiponectin, are inhibitors of osteoclastogenesis. Consistently, the total lack of adipocytes, including in the bone marrow, results in a complete absence of this adipokine, leading to the loss of its repressive activity in osteoclast differentiation.

AdipoRon is a non-peptidic agonist of adiponectin receptors (Okada-Iwabu et al., 2013). We thus perform the reverse experiment of the antibody blocking assay, adding AdipoRon to culture of bone marrow cells. Consistently, AdipoRon recapitulated the blocking effect of ACM on osteoclast differentiation irrespective of the origin of the osteoclast progenitors (bone marrow, spleen, or macrophage-enriched fraction) with IC50 in the micromolar range (Fig. 3a,b, supplementary Fig. S6. Furthermore, AdipoRon blocked osteoclast differentiation in *Pparg* deficient cells, confirming that intrinsic alterations do not play a role in the observed phenotype (Fig. 3c).

**Figure 3:**
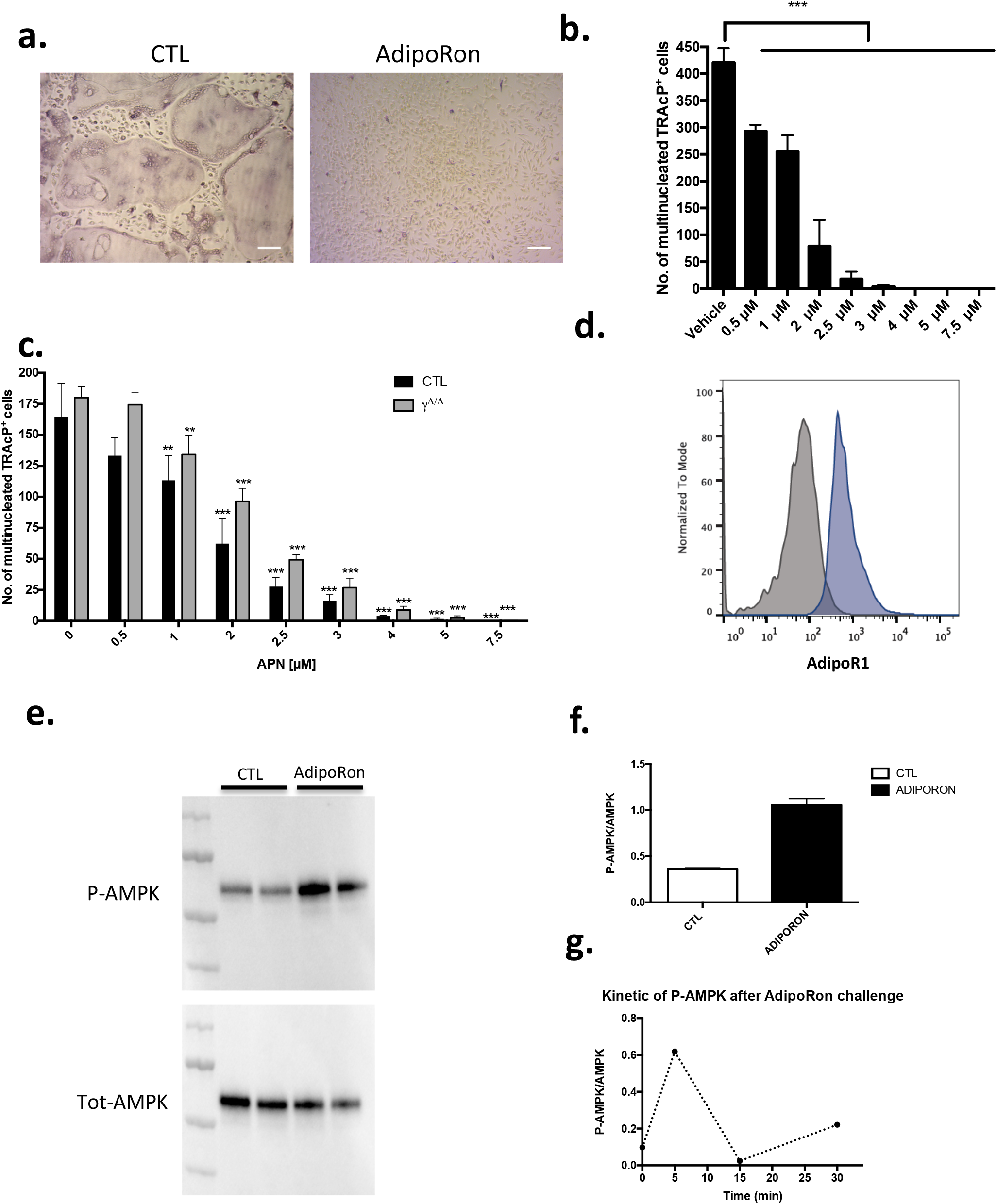
Adiponectin signalling activation blocks osteoclast differentiation. **a.** Representative images of TRAcP stained BM-derived osteoclasts differentiated in the presence or absence of AdipoRon (5μM). Scale bars=100μm. **b.** Dose response effect of AdipoRon on osteoclast differentiation using RAW264.7 cells (n=3). **c**. Effect of AdipoRon on osteoclast differentiation assay using bone marrow cells from *Pparg*^*Δ/Δ*^ (γ^*Δ/Δ*^) mice and control littermates (CTL). **d**. Expression of Type 1 Adiponectin Receptor (AdipoR1) at the surface of bone marrow derived osteoclasts demonstrated by flow cytometry. Grey curve represents unstained negative control; blue curve represents BM-derived osteoclasts. **e.** Western-blot analysis of sorted mature bone marrow-derived osteoclasts treated with AdipoRon (5μM). AdipoRon treatment activates AMPK phosphorylation (p-AMPK) in murine BM-derived osteoclasts (n=2 CTL; n=2 AdipoRon stimulated) **f**. Ratio of P-AMPK over total AMPK measured by densitometry **g**. Kinetic of AMPK phosphorylation (P-AMPK) after AdipoRon challenge in RAW264.7-derived osteoclasts. Data are presented as mean ± S.E.M. Statistical significance was determined by two-tailed unpaired *t*-test.

Previous studies demonstrated *in vivo* that mice treated with adiponectin expressing adenovirus have reduced osteoclast numbers and bone-resorption markers (Oshima et al., 2005). Using the macrophage cell line RAW264, prone to differentiate in osteoclasts, these authors further showed that adiponectin blocked *in vitro* osteoclastogenesis by inhibiting NFATc1. Blockade of NFATc1 was suggested to occur through activation of the AMPK signalling pathway, a key integrator of metabolic signals and target of adiponectin signalling (Yamaguchi et al., 2008). We thus tested whether this mechanism would apply *in vivo* and to primary cells. Flow cytometry analysis combining nuclei staining (Madel et al., 2018) and anti-AdipoR1 antibody, confirmed that the adiponectin receptor AdipoR1 is expressed also by mature osteoclasts generated from murine bone marrow cells (Fig. 3d). In agreement, western-blot analysis of AMPK phosphorylation confirmed that the Adiponectin signalling pathway is active in both precursors and mature osteoclasts sorted on the basis of their nuclei number (≥ 3) (Madel et al., 2018)(Fig. 3e-g).

AdipoRon was thus used to investigate the consequences of adiponectin receptor activation on murine and human osteoclasts. AICAR was used as a synthetic activator of AMPK. As seen in Figure 4a,b each of these molecules inhibited the formation of podosomes that are essential for bone resorption. Moreover, dorsomorphin, a reversible AMP-kinase inhibitor, could reverse the AdipoRon inhibitory effect (Fig. 4b). This is in line with the recently published effect of recombinant adiponectin on RAW264.7 cells, confirming the impact of adiponectin on both osteoclasts differentiation and podosome formation (Chen et al., 2018) and endorsing the use of AdipoRon as a surrogate activator for adiponectin receptors. Finally, adiponectin signalling regulates numerous cellular metabolic pathways including mitochondrial functions (Iwabu et al., 2010). AdipoRon treatment provoked disruption of the perinuclear mitochondrial network when added to culture of mature osteoclasts (Fig. 4c). Altogether, these results indicate that, beside its inhibiting action in osteoclast differentiation, adiponectin signalling exerts a pivotal role in mature osteoclast energy homeostasis regulation.

**Figure 4:**
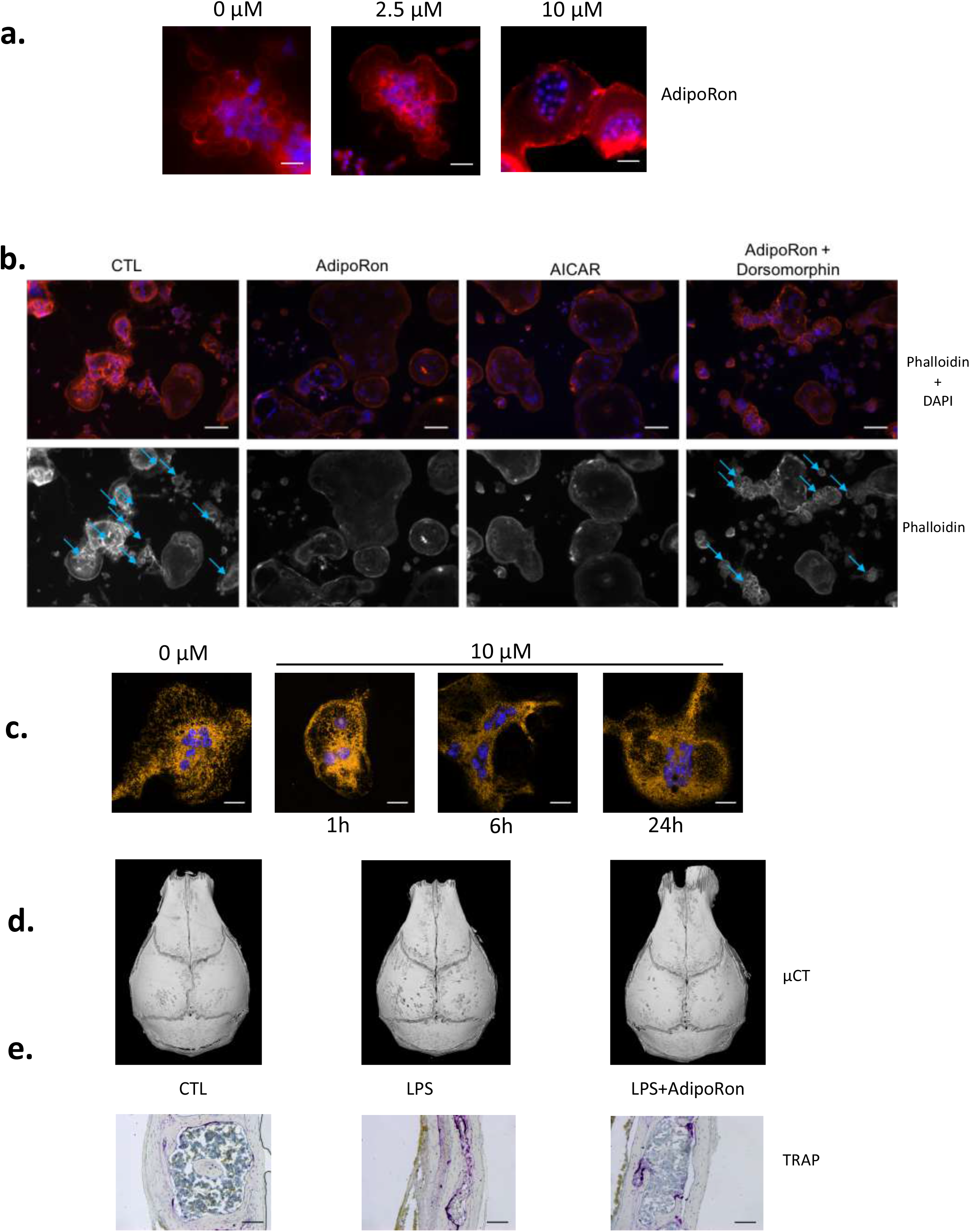
Adiponectin regulates OCL activity and podosome formation. **a.** Confocal microscopy of phalloidin-marked podosomes in RAW264.7-OCL in the presence of increasing concentrations of AdipoRon (representative images of n=3 per concentration) Scale bars=100 μm. **b.** Confocal microscopy of phalloidin-marked podosomes in the presence of AMPK modulating drugs (AdipoRon, 10μM; AICAR, 50nM; Dorsomorphin, 1.2μM) (representative image of n=3 per conditions white asterisks mark podosomes). Scale bars=50 μm. Blue arrows indicate podosome rings. **c**. Confocal microscopy of MitoTracker Orange in mature RAW264.7-OCL after AdipoRon (10μM) challenge at different time points. Scale bars=20 μm. **d**. Micro-CT of mouse skulls collected 3 days after mice received daily *s.c.* injections on the calvaria. Mice were injected with either 100 μg lipopolysaccharide from *Salmonella abortus equi* (LPS), or 100 μg LPS + 500 ng AdipoRon (LPS + AdipoRon) or vehicle (PBS) injection (control, CTL). (n=3 animals per group) **e**.TRAcP staining of mouse skull after LPS +/− AdipoRon challenge, (representative sections, n=3 animals per group). Scale bars=100 μm.

The inhibitory effect of AdipoRon was further tested *in vivo* on the well-described model of osteoclast bone resorption induced by lipopolysaccharide (LPS) injection into the mouse calvaria. Both micro-CT and TRAcP staining (Fig. 4 d,e) showed that AdipoRon administration resulted in a strong decrease in osteoclast numbers and bone resorption pits on calvaria.

We thus aimed to validate our findings on human cells and therefore investigated the impact of AdipoRon on hPBMC-derived osteoclasts. Consistent with the results obtained with mouse cells, low dose of AdipoRon significantly inhibited RANKL-induced human osteoclast differentiation (supplementary Fig. S7a,b). AdipoRon also activated AMPK phosphorylation in human PBMC-derived osteoclasts whithin minutes (supplementary Fig. S7c). At later time-point, i.e. within hours after the challenge with AdipoRon, the number of actin-containing podosome was drastically reduced in human osteoclasts (supplementary Fig. S7d). These original findings offer new therapeutic perspectives in inflammatory bone loss management by targeting adiponectin/AdipoR.

## Discussion

In the present report, using lipodystrophic mouse models, we evaluate how the total lack of adipose tissue may affect bone homeostasis. Taken together our results demonstrate that mature osteoclasts are sensitive to extrinsic adipose-derived metabolic signals such as adipokine and that bone marrow adiposity must be considered as a physiological regulator of bone homeostasis. Consistently with the literature, we identified adiponectin as one main inhibitory signal for osteoclastogenesis. Herein, we further show that AdipoRon, a pharmacological agonist of adiponectin receptors, inhibits osteoclast differentiation of precursors from different origins (bone-marrow, spleen and monocytic fraction). We also demonstrate that adiponectin signaling inhibits human osteoclast differentiation as well as activity of human mature osteoclast and explore how AMPK signaling act on mature osteoclast activity via podosome formation blockade. These original findings reveal a conserved adiponectin-mediated metabolic control of osteoclast activity and are clinically promising, in particular with regard to the treatment and prevention of excessive pathological bone-resorption conditions, such as in osteoporotic patients.

The first model used in the present study, the PPARγ null mice, is bringing up some decisive arguments with respect to the controversy on the role of PPARγ in osteoclastogenesis. Whereas initial studies reported that PPARγ is a key factor for osteoclastogenesis (Wan et al., 2007), they could not confirm such activity and proposed that only pharmacological activation of PPARγ had impact on osteoclastogenesis [12,23]. Herein, we definitely demonstrate that osteoclastogenesis can occur in the absence of PPARγ signal, and that other adipose tissue-derived signals, such as adipokines, affect osteoclast differentiation and activity. In contrast, the increase trabecular bone formation seen is vertebrae, in femoral epiphyses and distal diaphysis of *Pparg*^*Δ/Δ*^ mouse is consistent with the known role of PPARγ in inhibiting osteogenesis.

Using three complementary genetic models (2 distinct conditional null mice of PPARγ and the AZIP^tg/+^ mice, a PPARγ-independent lipoatrophy model), we show that the dramatic cortical bone alteration due to an excessive osteoclastogenesis in mice is linked to the lack of adipocyte and adipocyte-derived signals. The systemic and local activities of adipokines such as leptin and adiponectin have been recently reviewed (Cornish et al., 2018). Although leptin can act as a positive regulator of osteogenesis in *in vitro* systems, other studies in mice show that systemic leptin acting via the central nervous system is a potent inhibitor of bone formation (Ducy et al., 2000). Systemic adiponectin was reported to promote bone formation through its depressive action on the sympathetic tone in mice in age-dependent manner (Kajimura et al., 2013). However, *in vitro* differentiation assay showed that adiponectin inhibits osteoclast formation (Oshima et al., 2005; Yamaguchi et al., 2008). The observations *in vivo* are more controversial. Adiponectin-null mice exhibit an increased trabecular volume and fewer osteoclasts according to Wang *et al*. (Wang et al., 2014), whereas Shinoda *et al*. (Shinoda et al., 2006) found normal bone development in these mice. In contrast, Naot *et al.* (Naot et al., 2016) described a decreased cortical thickness in adiponectin-null female mice and Yang *et al*. (Yang et al., 2019) a decreased bone mass. Interestingly, the female bias of the phenotype observed herein was also found in adiponectin-null mice. Thus, adiponectin may have different effects on bone remodelling, depending on the activated autocrine, paracrine or systemic pathways (Tu et al., 2011). Our results from the osteoclastogenesis assay using bone marrow-derived cells and adiponectin-specific blocking antibodies support the important paracrine role of adiponectin in the inhibition of osteoclastogenesis in the normal BM environment. Direct paracrine activity is also consistent with the presence of the adiponectin receptors, AdipoR1 and AdipoR2, on the surface of differentiating osteoclasts (Pacheco-Pantoja et al., 2013). Nonetheless, we cannot exclude that other adipokines may also contribute to the paracrine effects of adipocytes on osteoclasts.

Identifying the source of adiponectin in the vicinity of osteoclast precursors and/or mature osteoclasts in bone marrow and bone compartment is challenging. Indeed, levels of adiponectin production by BM adipocytes remain controversial. Cawthorn et al. (Cawthorn et al., 2014) reported that BM adipocytes serve as a major source of circulating adiponectin exerting systemic effects during caloric restriction whereas others groups reported that bone marrow adipocytes express lower levels of adiponectin than adipocytes in adipose tissue (Li et al., 2019; Liu et al., 2011; Poloni et al., 2013). Others sources of bone marrow adiponectin have been recently described. Bone marrow PDGFRβ^+^VCAM-1^+^ stromal cells have been identified as major producers of adiponectin in the bone marrow (Mukohira et al., 2019). Single cell transcriptomic profiling of the mouse bone marrow stromal compartment indicated that LepR^+^ bone marrow mesenchymal stromal cells, which form a functional continuum with osteolineage cells, are expressing adiponectin (Baryawno et al., 2019). Beside their proliferative action on HSC and antiproliferative effects on myelomonocytic and early B lymphoid cells, these adiponectin-producing cells appears to be essential to the hematopoietic niche. In this context, our finding that adiponectin regulates osteoclast activity and differentiation in bone further strengthens the argument that a metabolic pathway driven by adiponectin controls many osteoimmunology processes, including remodeling of bone marrow hematopoietic stem cell niche, with potential implications for multiple human disease states including cancer.

The distinct behaviour and response of trabecular bone versus cortical bone is particularly highlighted in *Pparg*^*Δ/Δ*^ mice. Such contrasting phenotype have been previously observed. For example, increased or decreased mechanical strain (Ausk et al., 2013) or fluoride treatment increase the density of trabecular bone but decrease the density of cortical bone (Connett, Fluoride Action network, April 2012, http://fluoridealert.org/studies/bone03/). This bivalent phenotype is reminiscent of that observed when osteoblast and osteocyte apoptosis is prevented via targeted deletion of the proapoptotic proteins Bak/Bax (Jilka et al., 2014). In this context, a severe increase in cortical porosity was accompanied by increased osteoclast formation. The prolonged lifespan of osteoblasts was proposed by the authors to trigger diametrically opposed biological effects on bone homeostasis with an exaggeration of the adverse effects of ageing due to long-lived osteocytes (Jilka et al., 2014). In particular, osteocytes may increase intracortical remodelling via their secretion of RANKL and the subsequent activation of osteoclasts (Fuller et al., 1998; Jilka et al., 2014; Rinotas et al., 2014). In *Pparg*^*Δ/Δ*^ mice, the levels of *Sost* and *Keratocan* gene expression as markers of osteocytes and osteoblasts, respectively, are unchanged with respect to those found in their control littermates, thus refuting the possibility of an increased number of both cell types. In parallel, the increased expression of *Ctsk* and *Mmp9* likely contributes to the loss of cortical bone owing to high osteoclast activity, whereas the well-established role of PPARγ deficiency in bone formation via osteoblast activation is particularly highlighted by the increased trabecular bone density. Finally, it has been recently shown that osteoclasts have some site-dependent specificities and heterogeneity (Madel et al., 2019), that could contribute to the distinct behaviour of the cortical bone vs trabecular bone.

The difference in the intensity of the cortical porosity observed between the three lipodystrophic models we used may stem from different causes. On the one hand, the fact that the phenotype is more marked in the *Pparg*^*Δ/Δ*^ mice indicates that the expression of PPARγ in other cell types including osteoblasts and osteoclasts may dampen the phenotype in the AZIP and in the PPARγ^adipoKO^ models, irrespectively of adiponectin signals. However, a clear contribution of adipocytes remains highlighted by the results in AZIP mice which expressed functional PPARγ in all cell types. On the other hand, adiponectin gene expression, barely detectable in *Pparg*^*Δ/Δ*^ mice, is still present in the AZIP model and could contribute to the difference of phenotype penetrance between these two mouse models.

One important finding of the present report is the mechanism by which AdipoR1 signaling affects osteoclast activity. Indeed, here we demonstrated that AMPK was activated following AdipoRon challenge, which resulted in alteration of osteoclast metabolism, as exemplified by the alteration of the perinuclear mitochondrial network. We cannot exclude that direct or indirect signaling events may occur following adipoRon challenge that could interfere with osteoclast activity genetic program, such as Akt or NFATc1 signaling pathway as previously demonstrated with recombinant adiponectin (Tu et al., 2011; Yamaguchi et al., 2008). However, AMPK is central in this process, since its activation induced mitochondrial network alteration and podosome formation blockade, whereas AMPK blockade reverted these effects. Similarly, we also demonstrate that adiponectin signaling inhibits human osteoclast differentiation as well as activity of human mature osteoclast.

Altogether, these original findings reveal a conserved adiponectin-mediated metabolic control of osteoclast activity and are clinically promising, in particular with regard to the treatment and prevention of excessive pathological bone-resorption conditions, such as in osteoporotic patients. The present study implies that preserving functional adiponectin sensitivity in patients suffering from metabolic diseases (e.g. diabetes, obesity, metabolic syndrome) might be also beneficial for preventing bone fragility.

## Supplementary data

**Video 1:** Video of 3D micro-CT reconstruction of genetically induced lipoatrophic mice showing alterations of both trabecular and cortical femoral bone.

**Supplementary Table 1:**
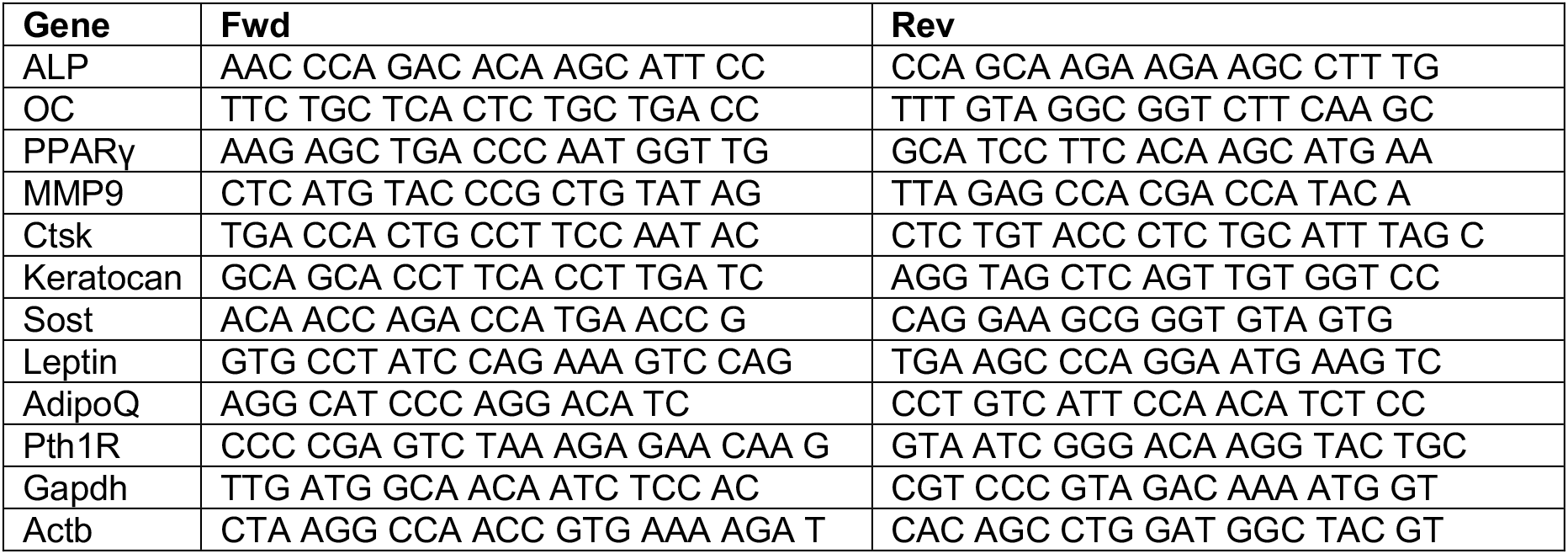
Primers used for q-RT-PCR.

**Supplementary Figure S1: Epiblastic PPARγ-deletion induces a more severe bone phenotype in female than in male. a.** Representative 3-D micro-CT of whole femur of male and female *Pparg*^*Δ/Δ*^ (γ^*Δ/Δ*^) mice. **b**. Representative 3-D reconstruction of trabecular bone in femoral epiphysis from female CTL, male and female *Pparg*^*Δ/Δ*^ (γ^*Δ/Δ*^) mice.

**Supplementary Figure S2: Osteoblastic activity is increased in mice with epiblastic deletion of PPARγ. a.** Representative 3-D micro-CT of the fourth lumbar vertebrae of CTL and *Pparg*^*Δ/Δ*^ (γ^*Δ/Δ*^) mice. n = 5 WT and 5 γ^*Δ/Δ*^. **b.** Trabecular bone volume fraction measured by micro-CT analysis of vertebra from γ^*Δ/Δ*^ (n=5) and control (CTL; n=5) littermates. **c.** Trabecular connectivity density measured by micro-CT analysis of vertebra from γ^*Δ/Δ*^ (n=5) and control (CTL; n=5) littermates. **d.** *Keratocan* and *Sost* mRNA levels in long bones of CTL (n=6) and γ^*Δ/Δ*^ (n=6) mice.

**Supplementary Figure S3: Increased cortical porosity in two lipodystrophic mouse models: A-ZIP^tg/+^ mice and mice with fat-specific deletion of PPARγ. a**. Representative 3D sections of midshaft femur from γ^adipoKO^ (n=3) and CTL (n=3) littermates. **b.** Femoral cortical porosity measured by micro-CT analysis of femur from γ^adipoKO^ (n=3) and control (CTL; n=3) littermates. **c.** Representative images of TRAcP staining of longitudinal sections of decalcified femurs, at mid-shaft levels, from control (CTL) or γ^adipoKO^ mice. CB: cortical bone; BM: bone marrow.

**Supplementary Figure S4: Parathyroid hormone (PTH) signalling**

The serum levels of PTH were assessed by Elisa Assay in 1-year-old male and female mice (*Pparg*^*Δ/Δ*^, AZIP^tg/+^, and their control littermates). Six males and six females were included per genotype. As there were no statistically significant differences between males and females, the analysis was performed using the total number of mice (mean ± SD). Lower panels: mRNA expression levels of Pth1r in the bones and in the kidneys of CTL *vs Pparg*^*Δ/Δ*^ mice.

**Supplementary Figure S5: Adiponectin expression levels in long bone and in adipocyte-conditioned medium**

**a.** RT-qPCR analysis of *Leptin* and *AdipoQ* mRNA levels in long bone from A-ZIP^tg/+^(n=8) and their CTL littermates (n=8) and from γ^*Δ/Δ*^ (n=7) and their CTL littermates (n=6). Data are presented as mean ± S.E.M. Statistical significance was determined by two-tailed unpaired *t*-test. **b**. Western-blot analysis of conditioned medium from undifferentiated (CTL) and differentiated 3T3-L1 adipocyte (A-CM) against adiponectin. Recombinant mouse adiponectin (200ng rmAdiponectin) was added to demonstrate specificity of the signal.

**Supplementary Figure S6: AdipoRon inhibits osteoclast (OCL) differentiation of multiple origins.**

*Left panels*: representative images of TRAcP stained (**a**.) BM-derived, (**b**.) spleen derived and (**c**.) macrophages enriched fraction osteoclasts differentiated in the presence or absence of AdipoRon (5μM). *Right panels*: the mean IC50 of AdipoRon inhibition of OCL differentiation was determined from the curve with the error of the fit (S.E.M.) Data are mean ± S.E.M. (n = 3 biological replicates).

**Supplementary Figure S7: AdipoRon inhibits human osteoclast (OCL) differentiation and activity through AMPK activation. a.** Dose response effect of AdipoRon on osteoclast differentiation from hPBMC (n=5). **b.** Representative images of TRAcP staining of hPBMC derived osteoclast differentiation in the presence or absence of AdipoRon (5μM). **c.** Western-blot analysis of hPBMC-derived osteoclasts treated with AdipoRon (5μM). AdipoRon treatment activates AMPK phosphorylation (p-AMPK) in hPBMC after 5 min. **d.** Fluorescence microscopy of phalloidin-marked podosomes in hPBMC derived OCL in the presence of AdipoRon (5μM, 12h) (representative images of n=5).

## Material & Methods

### Key resources table

| REAGENT or RESOURCE | SOURCE | IDENTIFIER |
| --- | --- | --- |
| Antibodies |  |  |
| Neutralizing adiponectin receptor 1 polyclonal antibody | Enzo Life Science | ALX-210-645-C200 |
| Adiponectin rabbit monoclonal antibody | Cell Signaling Technology | #2789 |
| AMPKα Rabbit mAb (D63G4) | Cell Signaling Technology | #5832 |
| Phospho-AMPKα (Thr172) (40H9) Rabbit mAb | Cell Signaling Technology | #2535 |
| Chemicals, Peptides, and Recombinant Proteins |  |  |
| Fetal Bovine Serum | GIBCO | 10082139 |
| Alpha MEM with NTP, dNTP | GIBCO | 22571 |
| AdipoRon | SIGMA | SML0998 |
| mRANKL | R&D systems | 462-TEC |
| hRANKL | R&D systems | 6449-TEC |
| Critical Commercial Assays |  |  |
| Acid Phosphatase, Leukocyte (TRAP) assay | SIGMA | 387A |
| murine serum CTX ELISA | LSBio | F21349 |
| murine serum OCN ELISA | LSBio | F12227 |
| murine serum PTH ELISA | LSBio | F23085 |
| Experimental Models: Cell Lines |  |  |
| 3T3-L1 pre adipose murine cell line | ATCC | CL-173 |
| RAW 264.7 murine macrophage | ATCC | TIB-71 |
| Experimental Models: Organisms/Strains |  |  |
| Sox2-Cre mouse | The Jackson Laboratory | 004783 |
| Adipoq-Cre mouse | The Jackson Laboratory | 010803 |
| PPARγ L2/L2 mouse | B Desvergne | PMID:15070754 |
| AZIP/F1 tg mouse | C Vinson | PMID:9784492 |
| Oligonucleotides (primers): see supplementary Table 1 |  |  |
| Software and Algorithms |  |  |
| CTan CTVol | Skyscan |  |
| ImageJ | Fiji |  |

### Mouse models

Genotype denomination follows the rules recommended by the Mouse Genome Database (MGD) Nomenclature Committee. Animal procedures were authorized by the Canton of Vaud veterinary service. Sox2CRE transgenic mice (SOX2CRE^tg/+^; Tg(Sox2-cre)1Amc/J) and AdipoQCRE (*Adipoq-Cre*^tg/+^) from The Jackson Laboratory, were maintained in the University of Lausanne Animal Facility. The series of matings allowing the generation of the SOX2CRE^tg/+^*Pparg*^*em**Δ/Δ***^ mouse (hereafter called *Pparg*^*Δ/Δ*^), which expressed the SOX2CRE transgene and have no *Pparg* functional alleles, as well as their control littermates (CTL) that have no SOX2CRE transgene and two functional *Pparg* alleles (*Pparg^fl/+^*) have been described in (Gilardi et al., 2019). The resulting genetic background of these mice are a mix of C57Bl6 and SV129L. PPARγ^adipoKO^ and their control littermates were generated as described in (Wang et al., 2013), in a mixed genetic background C57Bl6 and SV129L. AZIP/F1 mice on FVB background (Tg(AZIP/F)1Vsn in MGD, herein called AZIP^tg/+^) and wild type FVB controls were a kind gift from Dr. Charles Vinson and were generated as previously described (Moitra et al., 1998).

### Primary cell culture

Mouse bone marrow cells were isolated by crushing entire long bones into DMEM/3% FCS (GIBCO). Bone fragments were removed by filtration through a 70 μm filter mesh. Splenocyte suspensions were obtained by mashing the spleen through sieves into DMEM/3% FCS, washing by centrifugation then filtering through 70 μm filter caps. Primary osteoblasts from neonatal calvaria of *Pparg*^*fl/+*^ and *Pparg*^*Δ/Δ*^ were harvested by sequential collagenase type II (3 mg/mL) digestions of individual calvarium from 2 to 3-day-old mice. After genotyping, calvaria cells of the same genotype were pooled and plated accordingly. The cells were incubated at 37° C with 5% CO2 and media was changed every two days until they reached 80% confluency. To assess proliferation, primary osteoblasts were plated in 24-well plates at a concentration of 10,000 cells/well in EMEM media (GIBCO) containing 10% FBS, 100 U/ml penicillin and 100 ug/ml streptomycin (GIBCO).

### Osteoclastogenesis Assays and TRAcP staining of murine and human primary cells

Erythrocytes were eliminated from murine splenocyte and BM preparation or human PBMC using ACK lysis buffer. Cells were cultured in complete αMEM with 40 ng/ml M-CSF (ProSpec-Tany TechnoGene, Rehovat, Israel) and l0 ng/ml RANKL (R&D Systems, MN). Tartrate-resistant acid phosphatase (TRAcP) staining was performed after 4-6 days, using the Acid Phosphatase, Leukocyte (TRAP) Kit according to the manufacturer’s instructions (Sigma). Mature osteoclasts were identified as multi-nucleate TRAcP^+^ cells using light microscopy. 3T3-L1 cells (ATCC) were grown to sub-confluence in DMEM/10% FBS. Cells were then cultured in adipogenic conditions (DMEM/10%FBS, 0.5 mM 3-isobutyl-1-methylxanthine, 1 μM dexamethasone (Sigma), and 10 μg/ml bovine insulin (Sigma)) for 2 weeks, after which the medium was changed back to DMEM/10% FBS. Adipocyte Conditioned Medium (A-CM) was collected after 24 hours exposure to differentiated cells. Similarly, conditioned control medium (CTL) was obtained from undifferentiated 3T3-L1 cells. A-CM and CTL were concentrated 5X using Centriprep^TM^ (Millipore). Conditioned medium from either differentiated or undifferentiated 3T3-L1 cells was added during osteoclastogenesis from multipotent progenitor cells. For neutralisation experiments, 50% CM was incubated with different amounts of anti-adiponectin antibody (Enzo Life Sciences) for 30 min at room temperature then added during osteoclastogenesis. TRAcP staining to evaluate osteoclast numbers was performed as mentioned above.

### Osteoclast differentiation and TRAcP staining of RAW 264.7 cells

For osteoclastogenesis, a total of 1×10^4^ RAW 264.7 (ATCC) cells were seeded per well on 24-well plates in 500 μl αMEM containing 5% FCS (Hyclone, GE Healthcare), 1% penicillin-streptomycin (Gibco) as well as 50 μM β-mercaptoethanol (Gibco) and 30 ng/ml murine RANKL (R&D Systems). To study the impact of adiporon on osteoclast differentiation, 10 μM AdipoRon (Sigma-Aldrich) were added to the culture medium. Tartrate-resistant acid phosphatase (TRAcP) activity was examined using the leucocyte acid phosphatase kit (Sigma-Aldrich) according to manufacturer’s specifications. TRAcP^+^ cells with ≥3 nuclei were considered as osteoclasts. In addition, 5×10^3^ RAW 264.7 cells were seeded per well in a Nunc Lab-Tek II 8-well Chamber Slide system (ThermoFisher Scientific) in a total of 300 μl osteoclast differentiation medium containing 30 ng/ml RANKL. Fully differentiated osteoclasts were treated with 10 μM AdipoRon for the indicated time points. FACS analysis as well as sorting of mature OCLs for subsequent Western-blot analyses was performed as described previously (Madel et al., 2018).

### Measurement of bone morphology and microarchitecture

Unless otherwise indicated, one-year old females of each genotype were used for bone analyses. High-resolution micro-computed tomography (UCT40, ScancoMedical AG, Bassdorf, Switzerland) was used to scan the femur and the 5^th^ lumber vertebral body. Cortical and trabecular bones were evaluated using isotropic 12μm voxels. For trabecular bones, we assessed bone volume fraction (BV/TV, %), trabecular thickness (Tb.Th, μm), trabecular number (Tb.N, mm^−1^), and trabecular separation (Tb.Sp, μm). For cortical midshaft femur, we measured the average total volume (TV, mm^3^), bone volume (BV, mm^3^) and average cortical thickness (μm). SkyScan software (CTAn and CTVol) was employed to calculate cortical bone porosity of femur using a dedicate method after acquisition using a Skyscan 1272 (SkyScan 1272, Bruker, Brussels, Belgium) at a resolution of 10 μm/pix, and a threshold set up at 100-255, finally visualized in 3D.

### Serum protein assays

Blood was obtained by cardiac puncture immediately after euthanasia. After clotting and centrifugation, serum was collected and stored at −80°C until use. Serum CTX (carboxy-terminal collagen crosslinks), osteocalcin (OCN), alkaline phosphatase (ALP) and parathyroid hormone (PTH) were measured by ELISA assays according to the manufacturer’s protocol (LifeSpan Biosciences, Inc., Seattle, WA).

### Calvaria injection

Nine 6-week old C57/BL6 male mice were purchased from Envigo and housed in the local animal facility under a 12h light/12h dark cycle. The animals were randomly divided into three experimental groups, each group comprising three mice. Mice received daily *s.c.* injections on the calvaria before analysis on the third day. Group 1 was injected with 100 μg lipopolysaccharide from *Salmonella abortus equi* (LPS, Sigma-Aldrich), group 2 with 100 μg LPS + 500 ng AdipoRon (Sigma-Aldrich) and experimental group 3 underwent vehicle (PBS) injection. Calvariae were collected and fixed overnight in 4% formaldehyde at 4°C and then stored in 70% EtOH. Subsequent micro-CT analysis were used to evaluate the resorbed area. Calvariae were decalcified in 10% EDTA, embedded in paraffin and 5-micron sections were cut and stained for tartrate-resistant acid phosphatase (TRAcP, Sigma-Aldrich).

### Quantitative RT-PCR

Total RNA was isolated from total long bones, including the crude BM, or primary cell cultures, using TRIzol LS (Invitrogen) and purified with the RNeasy kit (Qiagen). RNA quality was verified by microfluidic (Agilent 2100 Bioanalyzer) and concentration determined with a NanoDrop spectrophotometer (Wilmington). Total RNA (500 ng/μg) was reverse transcribed using Script^TM^ cDNA synthesis kit (Bio-Rad) according to the manufacturer’s instructions. Real time qPCR was performed with SYBR® Green PCR mastermix using an ABI PRISM® 7900 PCR machine (Applied Biosystems). Results were normalized to glyceraldehyde-3-phosphate dehydrogenase (*Gapdh*) or beta-actin beta (*Actb*). The primer sets are indicated in the supplementary table 1.

### Western blot analysis

Mature OCLs were sorted based on their nuclei number (≥ 3) following the protocol described in Madel *et al.* (Madel et al., 2018). This protocol ensures that further analyses (FACS and western-blot analysis) are made with a pure mature cell population, avoiding contaminant effects due to the important number of non-osteoclastic cells always present in classical culture, as reported (Madel et al., 2018). The sorted mature OCLs were washed twice with ice-cold PBS and scraped in 1X Laemmli buffer. Cells were disrupted by sonication and centrifuged at 3000 rpm for 10 min. Protein samples were analyzed by SDS/PAGE (gradient 4-15% acrylamide, Criterion, BioRad) and electroblotted onto 0.2% PVDF membrane using Transblot Turbo (BioRad). After 1 h in blocking buffer (Tris-buffered saline (TBS)–Tween 20 with 5% fat free dry milk), membranes were blotted overnight at 4°C with Rabbit mAb against AMPKα (D63G4) (dilution 1:1 000, Cell Signaling Technology) or Phospho-AMPKα (Thr172) (40H9) (dilution 1: 1000, Cell Signaling Technology), diluted in TBS–Tween with 5% BSA. After three washing steps with TBS–Tween 20, blot was incubated for 1 h at room temperature with anti-rabbit IgG conjugated with horseradish peroxidase diluted 1:2000 in blocking buffer. After four additional washing steps with TBS–Tween 20, protein bands were detected by chemiluminescence using Chemi Doc XRS+ (BioRad).

### Imaging

Podosomes were visualized by phalloidin staining (Sigma-Aldrich) following manufacturer’s instructions. After removing unbound phalloidin conjugates, cells were labelled with 1 μg/ml DAPI (Sigma-Aldrich). Microscopic imaging was performed using a fluorescence microscope (Zeiss Axio Observer D1). For mitochondrial analysis, live cells were labeled with 200 nM MitoTracker Orange (ThermoFisher) according to manufacturer’s recommendations. After staining, cells were fixed in 4% formaldehyde, labelled with 1 μg/ml DAPI and subjected to z-stack confocal microscopy analyses (Zeiss LSM 710 on an inverted Axio Observer.Z1 stand). Images were processed using Fiji/Image J software (Schindelin et al., 2012).

### Statistical analysis

The data were analyzed using one- or two-factor ANOVA for multiple comparisons as appropriate. Two-group comparisons were performed using the Student’s *t*-test. ns: non-significant, * p<0.05, ** p<0.01, *** p< 0.001, **** p<0.0001.

## Acknowledgements

This work was supported by the Etat de Vaud (BD and AW) the FNRS (BD), the Région Grand Est (J-YJ), the Fondation Arthritis (DM), the french PIA project Lorraine Université d’Excellence, reference ANR-15-IDEX-04-LUE (J-YJ and DM) and the Fondation pour la Recherche Médicale (FRM, ECO20160736019) (MBM).

## Authors contribution

HF designed the experimental plan, performed the experiments, analyzed the data, and contribute to the original draft preparation. MBM, DDP, MS, CW, AW, CP, and BT performed experiments, analyzed the data, and prepared the vizualisation. JYJ, FG, SF, NB and CBW contributed to the conceptualization, analyzed the data, and reviewed/edited the manuscript. BD and DM conceived the study, designed the experimental plan, supervised the study, analyzed the data, wrote the manuscript, and acquired financial support for the project. All authors read and edited the manuscript. BD and DM are the guarantors of this work and, as such, had full access to all the data in the study and take responsibility for the integrity of the data and the accuracy of the data analyses.

