## Supplementary figures and images for "Adipose-tissue derived signals control bone remodelling"

### Sup Figure 1

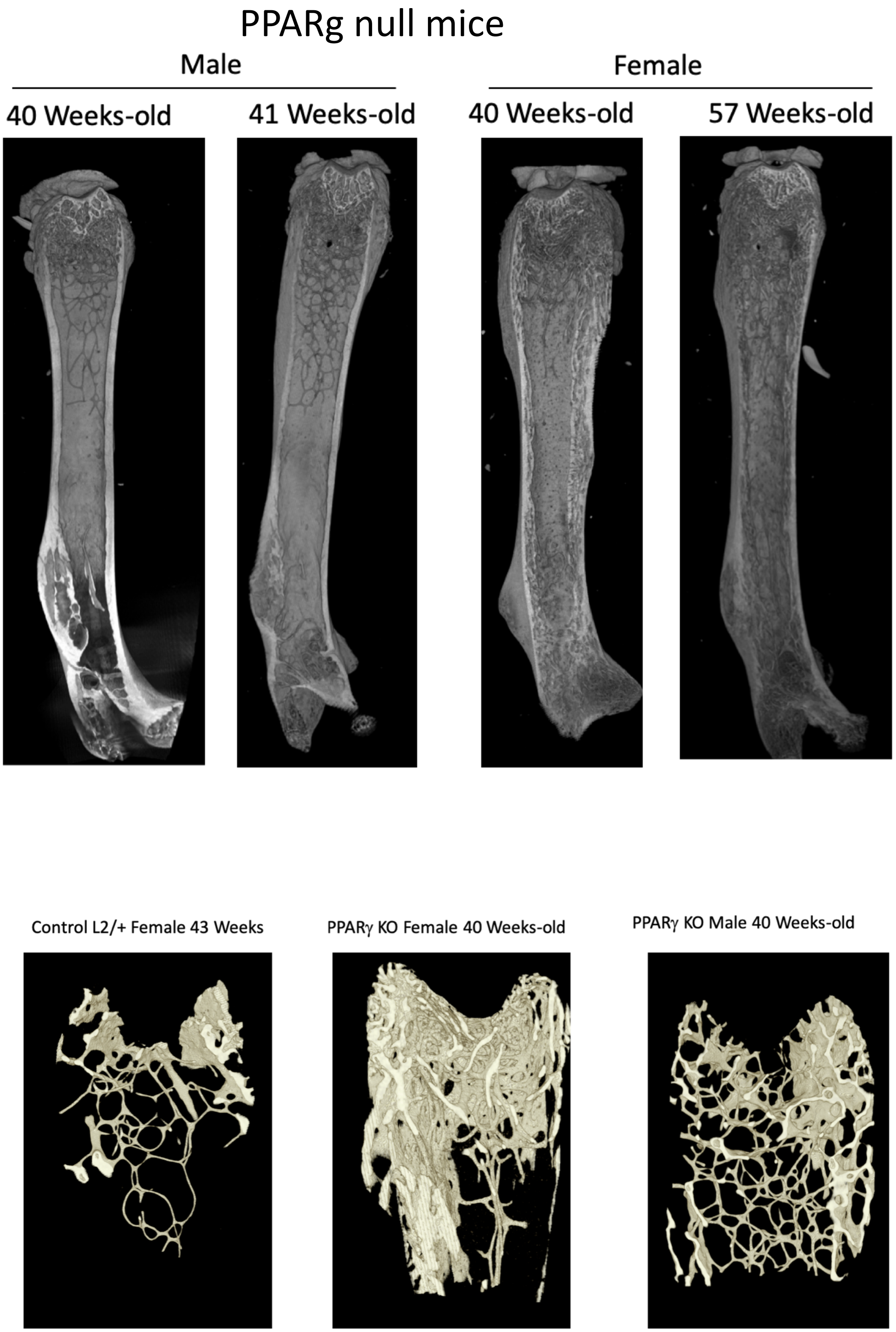

### Sup Figure 2

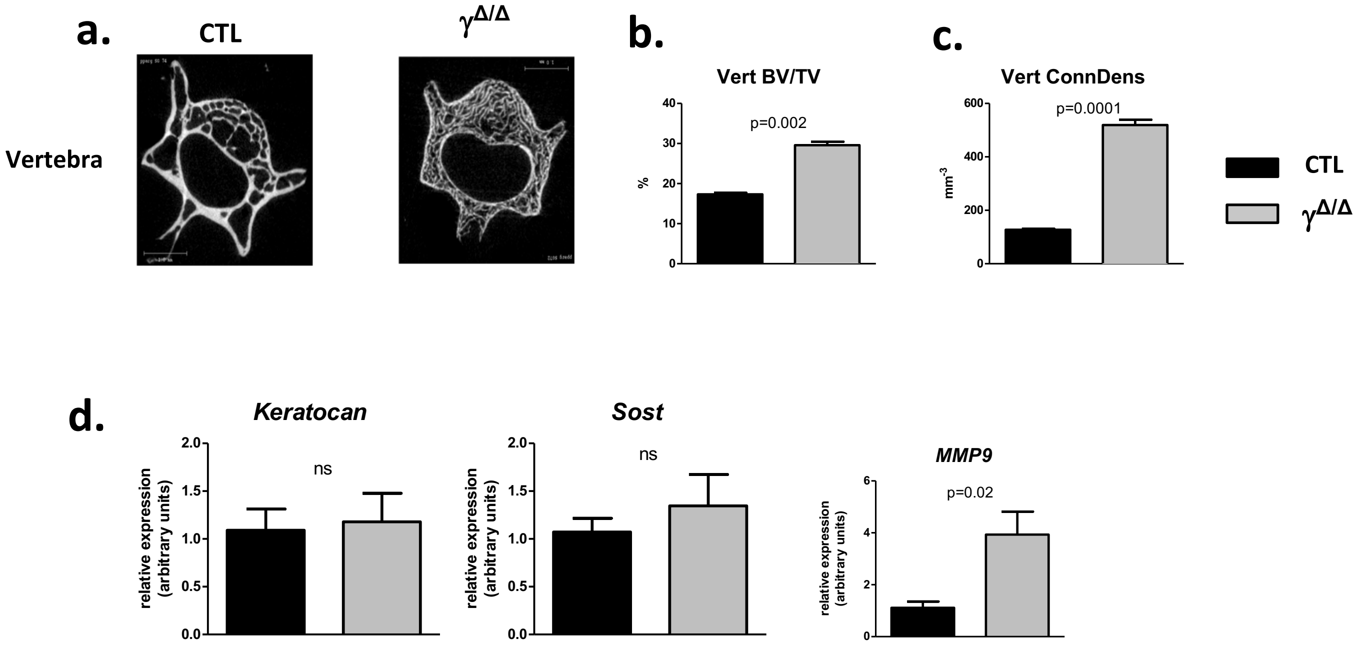

### Sup Figure 3

#### Figure S3

**a.**

**CTL** **$\gamma^{adipoKO}$**

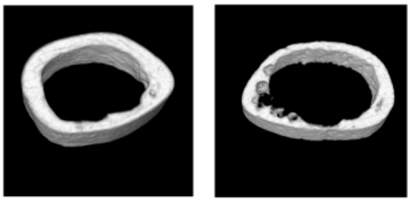

**b.**

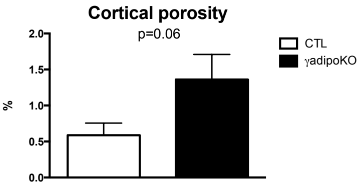


**c.**

**CTL** **$\gamma^{adipoKO}$**

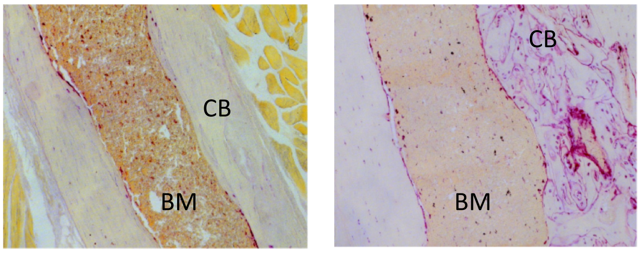

### Sup Figure 4

#### Figure S4 PTH signaling

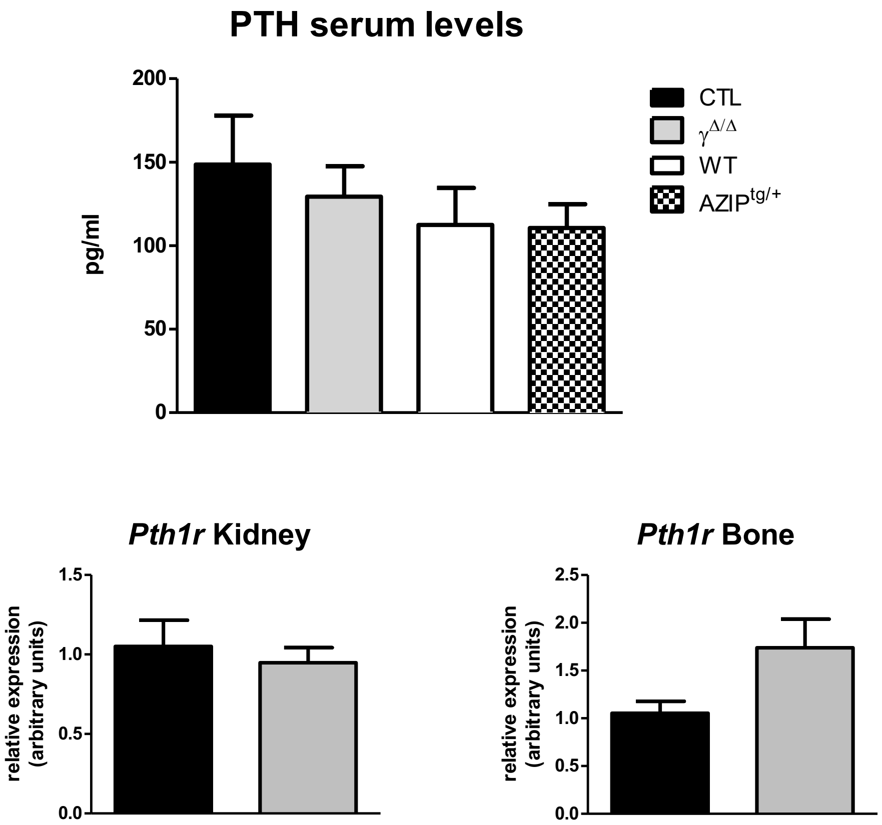

### Sup Figure 5

#### Figure S5

**a.**

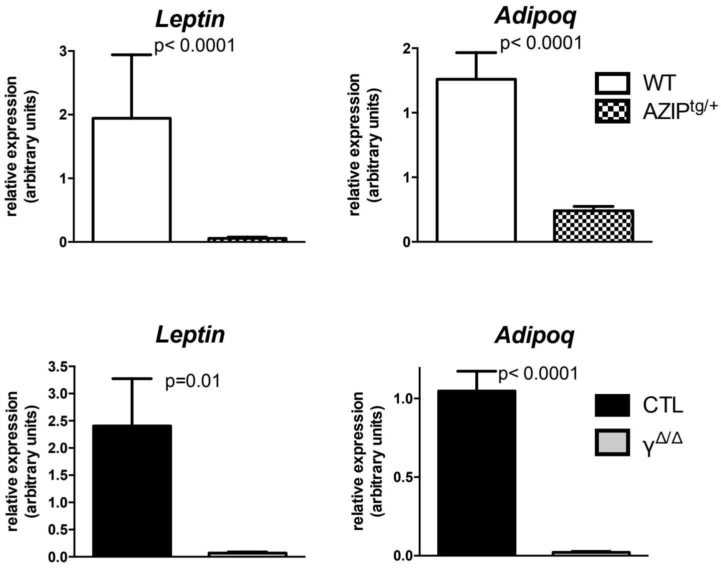


**b.** WB: Adiponectin

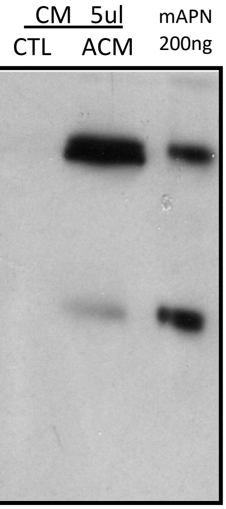

### Sup Figure 6

#### Figure S6

a.

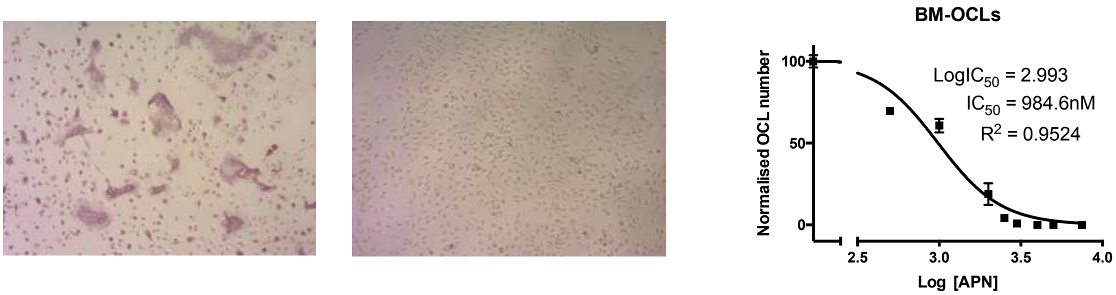


b.

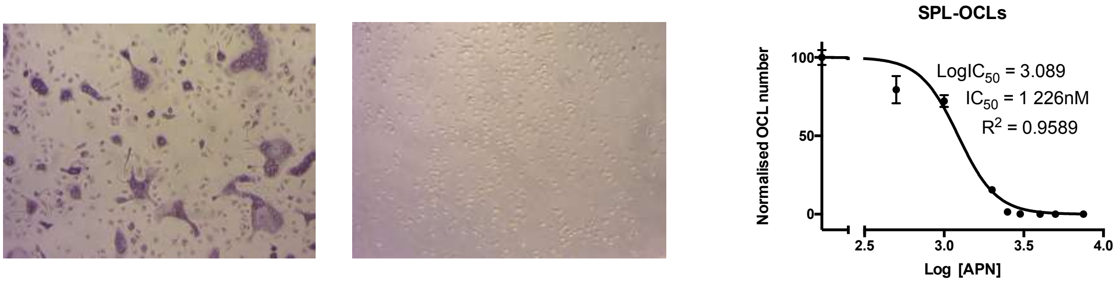


c.

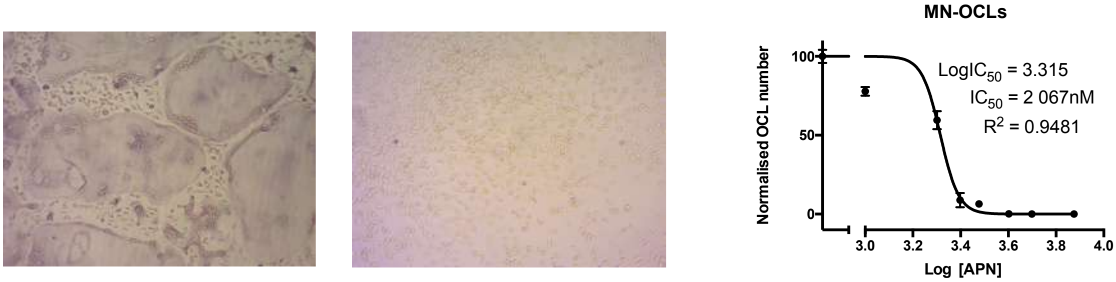

### Sup Figure 7

#### Figure S7

##### AdipoRon inhibits human OCL differentiation and activity

a.

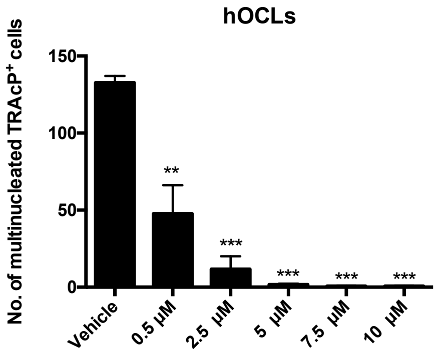


b.

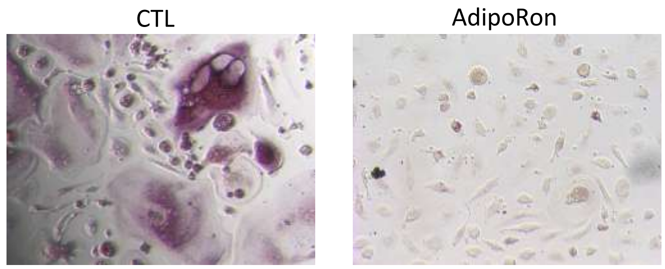


c.

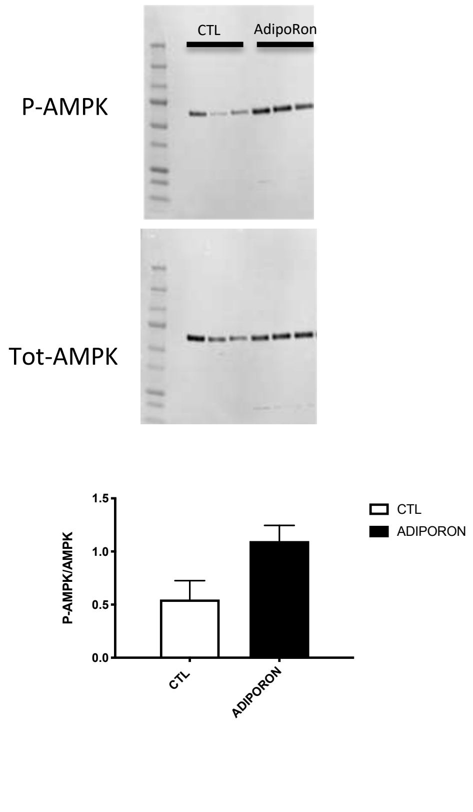


d.

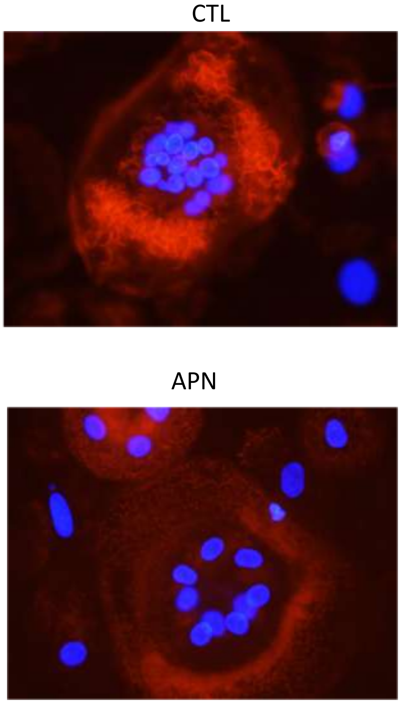
